# Influenza A Virus Negative Strand RNA is Translated for CD8+ T Cell Immunosurveillance

**DOI:** 10.1101/261966

**Authors:** Heather D. Hickman, Jacqueline W. Mays, James Gibbs, Ivan Kosik, Javier Magadan, Kazuyo Takeda, Suman Das, Glennys V. Reynoso, Barbara F. Ngudiankama, JiaJie Wei, John P. Shannon, Daniel McManus, Jonathan W. Yewdell

**Affiliations:** Laboratory of Viral Diseases, National Institutes of Allergy and Infectious Diseases, National Institutes of Health, Bethesda, MD 20892.; Microscopy and Imaging Core Facility, Center for Biologics Evaluation and Research, Food and Drug Administration, Silver Spring, MD 20993

**Keywords:** CD8+ T cells, immunosurveillance, influenza

## Abstract

To probe the limits of CD8+ T cell immunosurveillance, we inserted the model peptide SIINFEKL into influenza A virus (IAV) negative strand gene segments. Although IAV genomic RNA is widely considered as non-coding, there is a conserved, relatively long open reading frame present in the genomic strand of segment eight, encoding a potential protein termed NEG8. The biosynthesis of NEG8 from IAV has yet to be demonstrated. While we failed to detect NEG8 protein expression in IAV infected cells, cell surface K^b^-SIINFEKL complexes are generated when SIINFEKL is genetically appended to the predicted COOH-terminus of NEG8, as shown by activation of OT-I T cells *in vitro* and *in vivo*. Moreover, recombinant IAV encoding SIINFEKL embedded in the negative strand of the NA-stalk coding sequence also activates OT-I T cells *in vivo*. Together, our findings demonstrate both the translation of sequences on the negative strand of a single stranded RNA virus and its relevance anti-viral immunosurveillance.

**Significance:** Every gene encodes complementary information on the opposite strand that can potentially be used for immunosurveillance. In this study, we show that the influenza A virus “non-coding” strand translated into polypeptides during a viral infection of either cultured cells or mice that can be recognized by CD8+ T cells. Our findings raise the possibility that influenza virus uses its negative strand to generate proteins useful to the virus. More generally, it adds to a growing literature showing that immunosurveillance extends to gene sequences generally thought not to be converted into proteins. The relevance of translating this “dark” information extends from viral immunity to cancer immunotherapy and autoimmunity.

## Introduction

Influenza virus (IAV) causes significant worldwide morbidity and mortality due to its efficient evasion of humoral immune responses. IAV elicits a robust CD8+ T cell response, and extensive evidence in man and animals supports a role for CD8+ T cells in limiting viral replication and reducing morbidity and mortality in hosts whose antibody responses fail to prevent infection (1–4).

CD8+ T cell immunosurveillance encompasses oligopeptides encoded by each of the eight IAV gene segments. The exact peptides recognized by any infected individual is governed largely by their classical MHC class I genes (HLA-A,-B,-C in humans, H-2 K, D, L in mice) (5). In most species, each class I locus has thousands of alleles whose peptide specificity varies based on polymorphisms in and around the peptide binding groove.

Following viral infection, viral peptide ligands are generated extremely rapidly, with rates that parallel the translation of their source gene products (6–8). As viral proteins typically exhibit half-lives on the order of tens of h, the kinetic connection of peptide generation with protein synthesis, not degradation, strongly implies that viral peptides derive from a distinct pool of nascent gene products, termed DRiPs, for defective ribosomal products (9).

Though the biochemical nature of DRiPs generated from wild-type proteins remains largely undefined, viral peptides can clearly originate from non-canonical translation products (10). This includes downstream initiation on AUG codons (11), read through of stop codons (12, 13), frame shifting (14, 15), initiation on CUG or other alternative start codons (16), stop codon read through (12, 17) and translation of viral RNA in the nucleus (18). Inasmuch as each of these mechanisms would likely generate a defective gene product that is rapidly degraded, they provide a clear source of immunologically relevant DRiPs.

Negative strand viruses provide an intriguing opportunity for additional non-canonical translation in immunosurveillance, as the negative strand provides a potential source of peptides. Influenza viruses are of special interest since RNA transcription occurs in the nucleus, which may increase the possibility of non-canonical translation (18–21), or insertion of negative coding information into mRNA by RNA splicing, intended or otherwise. Adding to the interest, a large open reading frame (ORF) encoding 167 or more residues (depending on the viral strain) is present in the genomic strand of segment 8 (**Fig. 1A**), that has been conserved in human IAV isolates for the last 100 years (22, 23). MHC peptide prediction algorithms have identified several NEG8-derived peptides capable of eliciting T cell responses after IAV infection, and one peptide is reported to be immunogenic in IAV infected-mice (24). By contrast, evidence suggests that NEG8 is not under positive selection in nature among IAV isolates (25).

**Figure 1.**
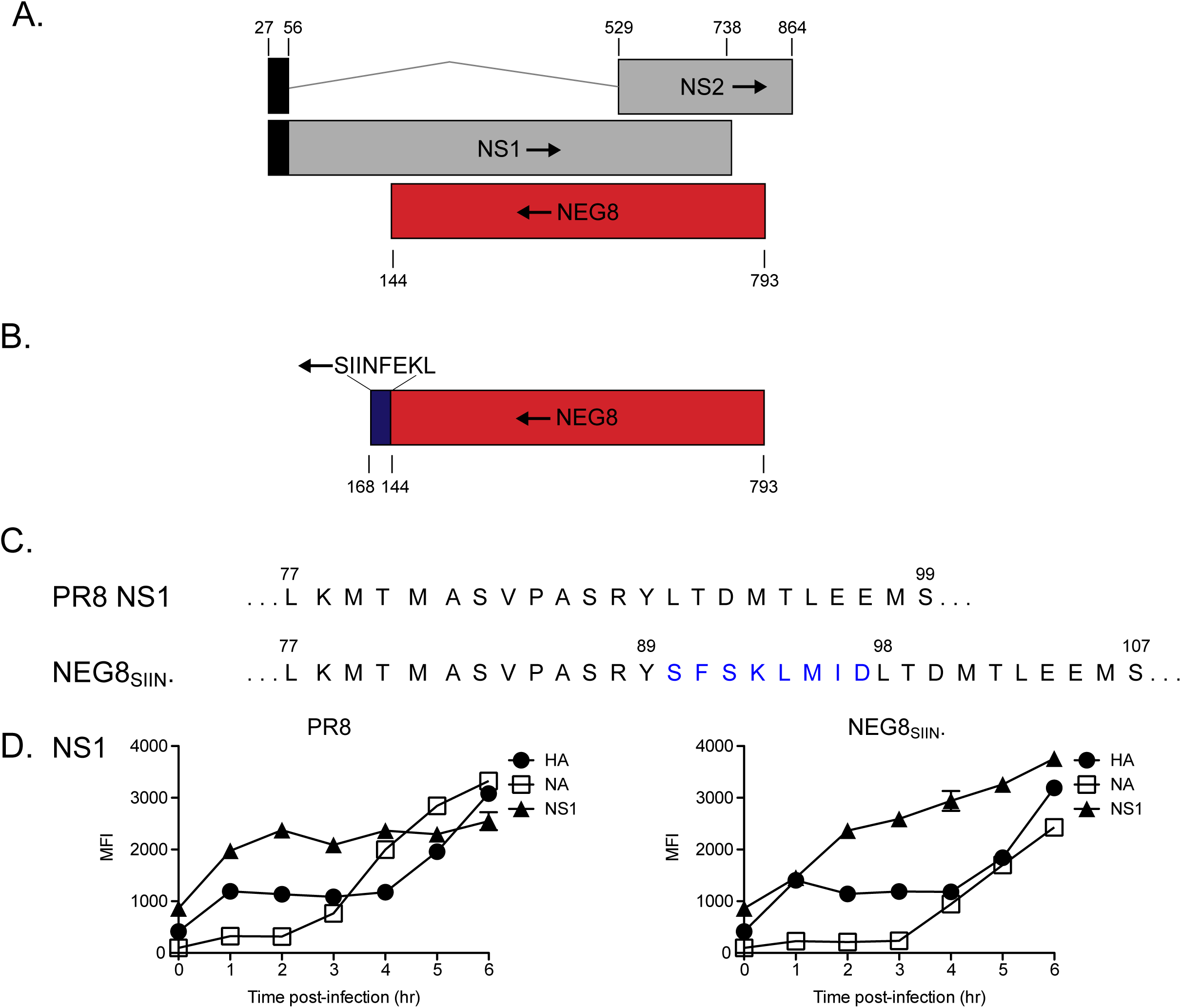
Production of recombinant virus expressing NEG8_SIIN_. A) NEG8 ORF on the negative, genomic strand of IAV segment 8 and its overlap with NS1/NS2. B) Strategy for NEG8SIIN production. C) Residues inserted into NS1 by insertion of negative-orientation SIINFEKL. D) Expression of viral proteins NA (open squares), HA (closed circles), and NS1 (closed triangles) after DC2.4 infection with PR8 (right) or NEG8SIIN. (left). Time=h. Error bars=means +/- SEMs.

Here we explore immunosurveillance of viral negative strand-encoded peptide by inserting the model peptide SIINFEKL into several locations in genomic “non-coding” RNA in IAV, including NEG8. SIINFEKL is a highly immunogenic peptide presented by H-2 K^b^ to CD8+ T cells (26), and its expression *in vitro* and *in vivo* can be monitored at high sensitivity using OT-I transgenic T cells (27). Our findings clearly show that immunosurveillance extends to genetic information encoded by the genomic strand of negative strand RNA viruses.

## Results

### NEG8 Encodes an ER-localizing Protein Undetected in IAV-Infected Cells *in vitro*

We initially explored IAV negative strand translation by attempting to demonstrate NEG8 translation, which has not been verified experimentally in the context of an IAV infection. We infected MDCK or HeLa cells with A/PR/8/34 (PR8) and used polyclonal rabbit antibodies raised to synthetic peptides corresponding to the predicted NH_2_- or COOH- termini in immunoblots of whole cell lysates or in immunofluorescence of fixed and permeabilized cells. Although rabbit sera demonstrated high titers against the immunizing peptides in ELISA, they failed to specifically stain PR8-infected cells in immunofluorescence, or to give a specific signal in immunoblots of total infected-cell lysates. We also failed to detect NEG8 in similar experiments using antibodies to epitope tags (His, Myc, Myc repeated 3X to increase anti-Myc Ab binding) that we had genetically appended to the predicted COOH-terminus of NEG8 in a PR8 recombinant virus.

By contrast, we could easily detect NEG8 with COOH-terminal GFP expressed from either a plasmid or a recombinant vaccinia virus (rVACV), consistent with a previous report that the protein is stably expressed by a recombinant baculovrius (28). As predicted (29), rVACV-expressed NEG8-GFP is present in the ER and post-ER compartments (Supplemental Figure 1), as shown by clear localization to the nuclear membrane and co-localization of GFP signal with Abs specific for ER (PDI) and trans-Golgi complex (TGN46). Reasoning that NEG8 might be metabolically unstable in IAV-infected cells, we treated PR8-infected cells with MG132 to inhibit proteasome-mediated degradation. This did not enable detection of PR8-encoded native or epitope tagged NEG8 via immunoblotting or immunofluorescence.

Thus, although NEG8 is clearly expressed from a plasmid or rVACV as a reasonably stable protein, we repeatedly failed to find evidence that it is expressed by IAV-infected cells in cultured cells.

### SIINFEKL Tagging of NEG8 Reveals its Translation in IAV-Infected Cells *in vitro*

Our failure to detect NEG8 protein could be due to a number of factors other than the complete absence of translation from genomic RNA. CD8+ T cells can provide an exquisitely sensitive measure of translation, and indeed, Zhong *et al.* reported that of 4 predicted K^b^/D^b^ binding peptides in the PR8 NEG8 ORF, one, corresponding to residues 33–40, binds to K^b^ and activates IAV-induced CD8+ T cells *in vitro* (24). We were unable to confirm that NEG833-40 binds K^b^ by a standard flow-based K^b^ stabilization assay (Supplemental Figure 2).

To further examine potential immunosurveillance of IAV genomic RNA we generated NEG8_SIIN_, a PR8-recombinant virus with SIINFEKL fused to the putative NEG8 COOH-terminus (**Fig. 1B**). SIINFEKL forms a highly stable complex with the mouse K^b^ class I molecule that can be detected by either the 25D-1.16 mAb, or at highest sensitivity, by OT-I transgenic T cells.

Inserting sequences into genomic RNA will, of course, alter proteins encoded by the positive-sense strand. Although PR8 NEG8_SIIN_ has an 8-residue insertion between resides 89 and 90 of NS1, the virus replicates in embryonated chicken eggs at levels similar to wild-type (*wt*) virus. Sequencing of egg-grown stocks confirmed the presence of the SIINFEKL-encoding insert, demonstrating that NS1 can function with an essentially random insertion of eight amino acids after residue 89. Though the function of NS1 with an 8-residue insertion might seem surprising, it is likely explained by the perhaps not completely fortuitous location (see Discussion) of the insertion in the extended polypeptide that links the NS1 effector and RNA binding domains (Supplemental Figure 3). Following infection of DC2.4 cells (a B6-mouse derived, dendritic cell-like cell line (30)) with NEG8_SIIN_ or *wt* PR8, flow cytometry revealed similar expression kinetics of NS1 using fixed and permeabilized cells and HA- or NA- staining prior to fixation (**Fig. 1D**).

We next incubated NEG8-SIIN- or PR8-infected DC2.4 cells with OT-I CD8+ T cells, and measured T cell activation after 1 d by induction of the CD69 or CD25 activation markers or by cell division (measured by dilution of CFSE label) (**Fig. 2**). Two positive controls, SIINFEKL peptide-supplemented cultures and cells infected with PR8 with SIINFEKL inserted into the NA stalk (PR8 NA_SIIN_) (31, 32), confirmed that DC2.4 expressing K^b^-SIINFEKL can robustly activate OT-I cells under these conditions. By contrast, uninfected cells or cells infected with PR8 lacking SIINFEKL failed to express activation markers or to divide, demonstrating the expected requirement for DC2.4 cell presentation of K^b^-SIINFEKL complexes. Remarkably, NEG8_SIIN_-infected cells activated OT-I cells at similar or even higher levels relative to the positive controls, as assessed by activation markers or cell division. To our knowledge, this provides the first direct evidence of translation of genetic information from the IAV negative strand.

**Figure 2.**
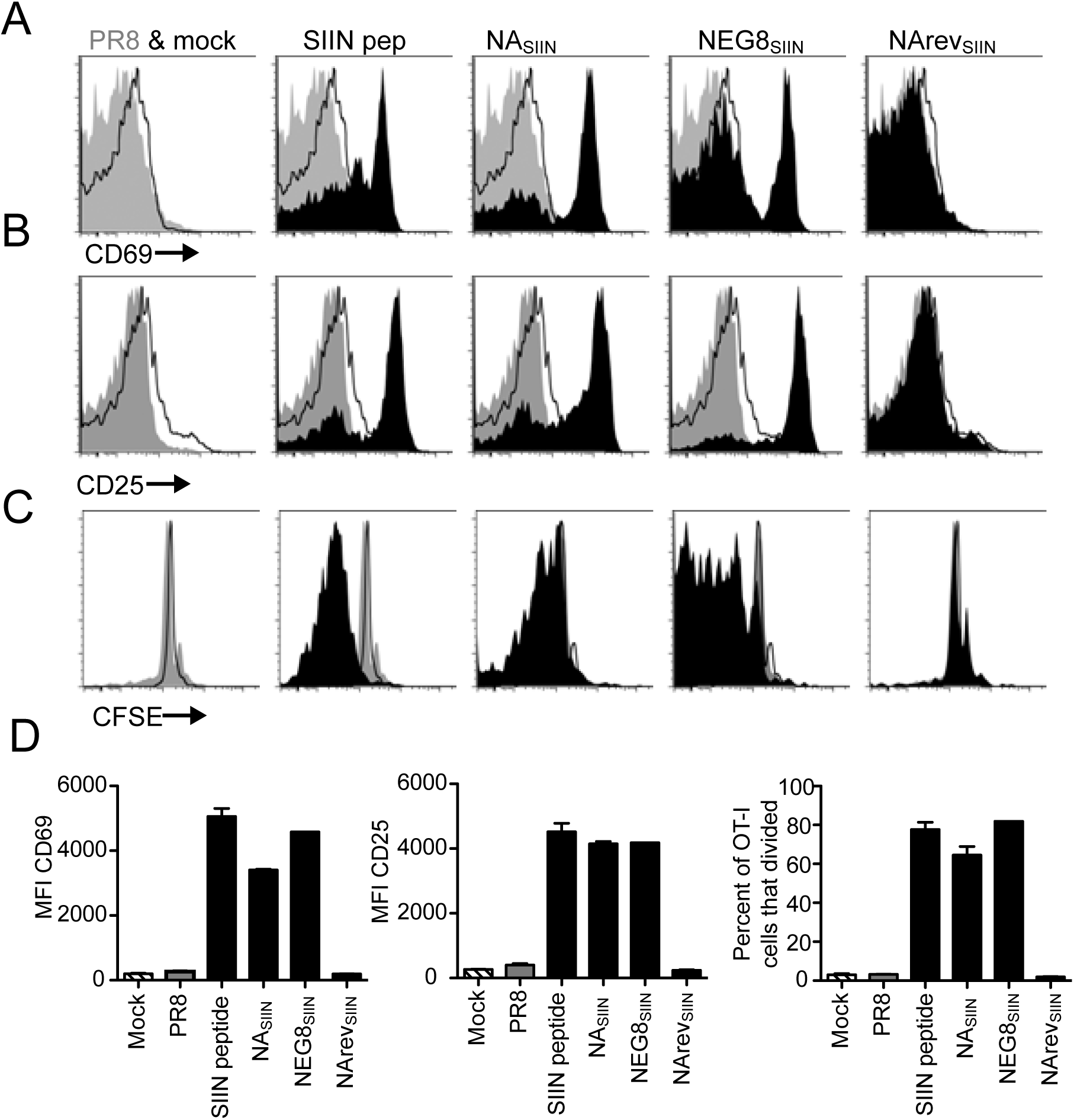
SIINFEKL is produced *in vitro* after NEG8_Siin_ infection and activates OT-I CD8^+^ T cells. A-B) CD69 (A) or CD25 (B) expression 24 h. after co-culture with infected cells. Mean fluorescence intensities (MFI), right. C) CFSE dilution (indicating cellular division) 48 h. after co-culture. Error bars=means +/- SEMs. Statistics=two-tailed student’s t test. Experiments were repeated twice with 2–3 replicates/group.

Generating additional recombinant PR8 viruses (**Fig. 3**), revealed that a start codon at the predicted NH_2_-terminus of NEG8_SIIN_ is not required for generation of SIINFEKL from infected DC2.4 cells, as determined by *in vitro* induction of CD69 on OT-I T cells using a virus in which the initiating Met is replaced by Thr (Met(-)NEG8_SIIN_). Further, a stop codon immediately upstream of SIINFEKL (StopNEG8_SIIN_) abrogates K^b^-SIINFEKL generation, though not completely, consistent with read-through translation. Note that flow cytometric detection of NA on live cell surfaces confirmed that cells were well infected by the various PR8 viruses used (all over 90% positive).

**Figure 3.**
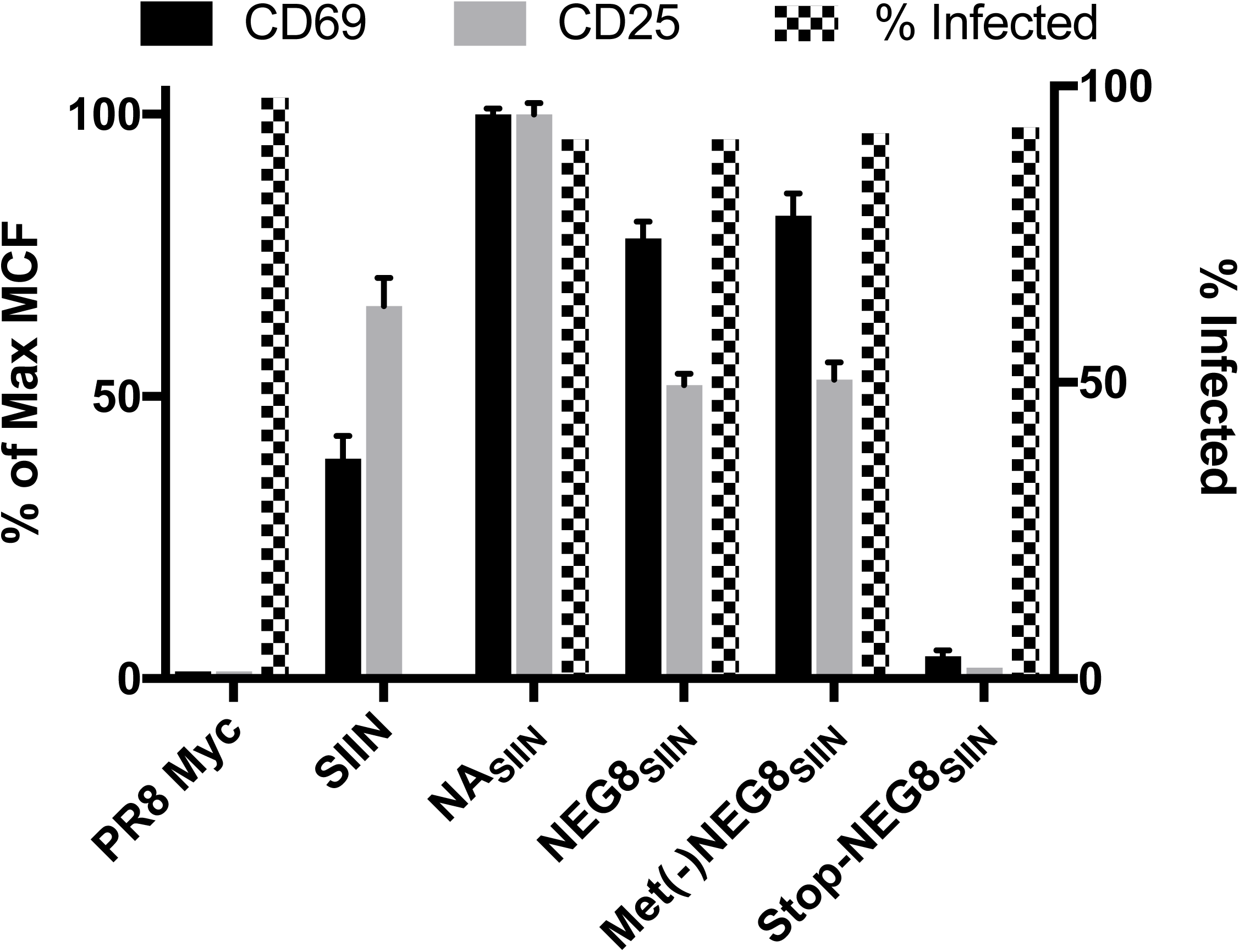
Characterization of NEG8_SIIN_ translation. DC2.4 cells were infected with the rPR8 viruses indicated and tested 8 h post-infection for NA expression (% infected) by flow cytometry using directly conjugated NA2-1C1 mAb. After infection, a separate aliquot of cells were co-incubated for 24 h with OT-I T cells, which were then stained for expression of CD69 and CD25 using directly conjugated mAbs. Data represent the MCF determined by flow cytometry, normalized to maximal expression, error bars represent the SEM. Two additional experiments gave highly similar results.

These findings indicate that initiation can occur downstream of the first AUG in the NEG8 ORF. ATGpr bioinformatic software (http://atgpr.dbcls.jp/) predicts that initiation is nearly equally likely to occur on the second of the 6 NEG8 AUG codons (reliability score of 0.14 vs. 0.12 for first vs. second AUG), which would encode a 93-residue polypeptide. Initiation could also occur on any of other 4 AUG codons, despite their low score (all at background levels), since the accuracy of the algorithm falls off at the bottom end, with a specificity of only 20%. In addition, we note the presence of 4 CUG codons that could support initiation, particularly since CUG initiation is likely favored during IAV infection (16, 33).

It important to mention that while PR8 NEG8_SIIN_-infected cells consistently activated OT-I CD8+ T cells, we only detected K^b^-SIINFEKL complexes on the cell surface using the K^b^-SIINFEKL complex-specific 25-D1.16 mAb (34) in aminor fraction experiments; always perilously close to the limits of detection. As OT-I cells are approximately 30-fold more sensitive than 25-D1.16-based detection (34), this is consistent with the generation of low amounts of K^b^-SIINFEKL from PR8 NEG8_SIIN_-infected cells.

### NEG8_SIIN_ is Sufficiently Translated *in vivo* to Enable Immunosurveillance

To establish the *in vivo* relevance of these findings, we transferred 2 × 10^5^ CD45.1^+^ OT-I CD8+ T cells into C57Bl/6 (CD45.2^+^) mice and infected them one day later via intraperitoneal injection with the various recombinant PR8 viruses (**Fig. 4**). Seven d p.i., we enumerated OT-I (CD45.1^+^) cells in spleens and peritoneal exudate cells (PECs) by flow cytometry. In control PR8-infected mice, there were few (< 0.2%) OT-I T cells present in the spleen or PECs. In contrast, PR8 NAsiin infection increased OT-I T cell percentages in both the spleen and PECs (20.8 ± 1.6 and 31.9 ± 2.5% or total cells, respectively). In NEG8_SIIN_-infected mice, 2.8 ± 0.2% of splenocytes and 5.5 ± 1.2% of PECs were OT-I cells, approximately 10–20 times greater than background. Notably, the total number of CD45.1^+^ OT-I T cells recovered was far greater than the number adoptively transferred, indicating OT-I proliferation in response to SIINFEKL rather than solely altered recruitment (**Fig. 4** **A**, **B**, far right panels).

**Figure 4.**
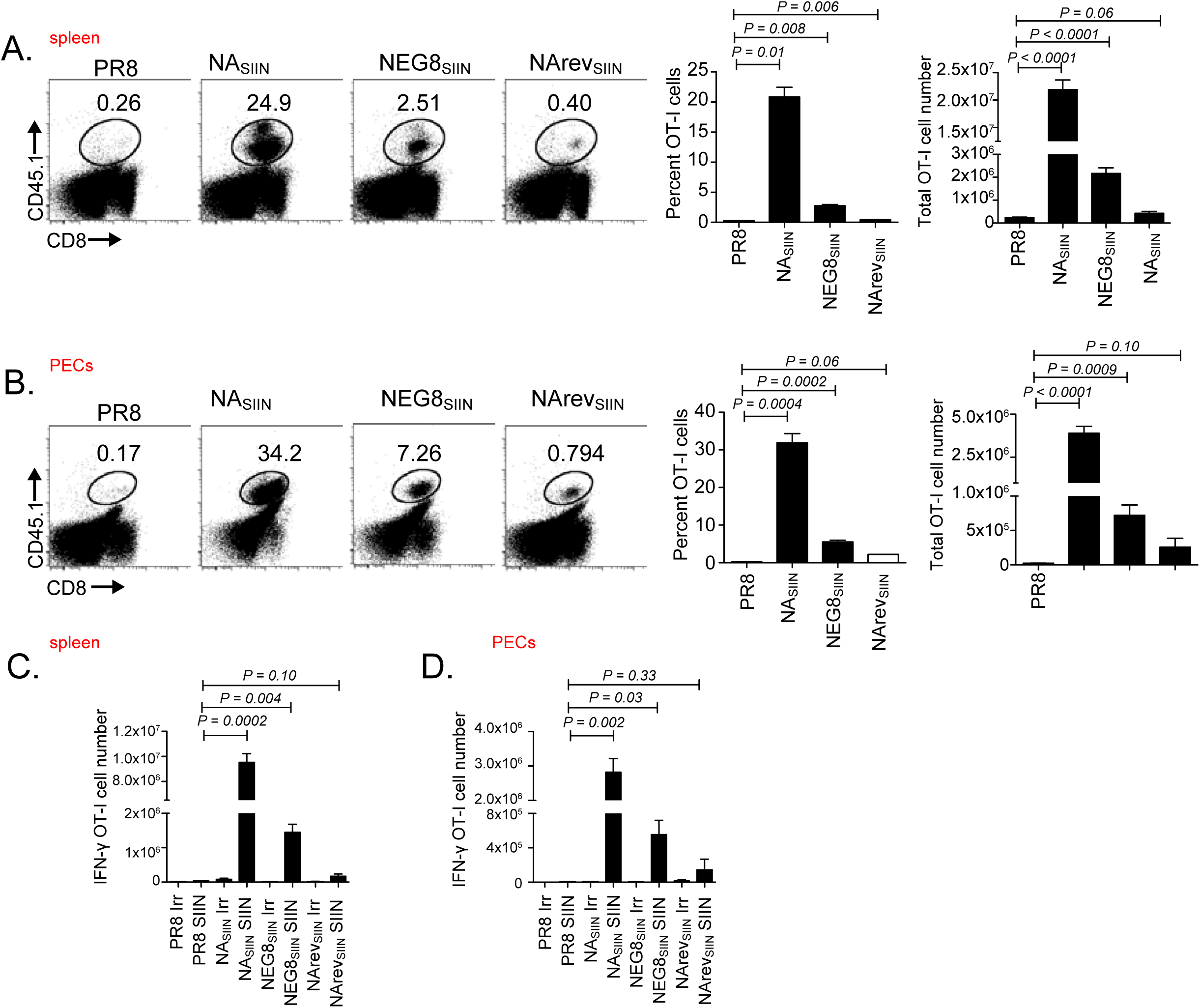
NEG8-SIIN is produced *in vivo* and recognized by the antiviral immune response. A) Flow cytometric dot plots showing gates for CD8^+^ CD45.1^+^ transferred OT-I cells in the spleen (A) or peritoneal exudate cells (PECs, B) 7 d.p.i. Means and SEMs for percent of cells (middle) and total cell number (right panels) are indicated. C-D) Numbers of OT-I T cells producing IFN-γ in the spleen (C) and PECs (D). Means, SEMs shown. Statistics = two-tailed *t* test. Experiment was performed twice with 3 mice/group.

To confirm OT-I activation by NEG8_SIIN_ we assayed splenic and PECs for IFN-γ synthesis by stimulating *ex vivo* with SIINFEKL or an irrelevant peptide (**Fig. 4C, D**). A majority of OT-I cells in either NA_SIIN_- or NEG8_SIIN_-infected mice produced IFN-γ in response to re-stimulation with SIINFEKL but not an irrelevant peptide, indicative of Ag-experienced, effector CD8^+^ T cells.

Taken together these data clearly demonstrate that NEG8_SIIN_ activates CD8+ T cells *in vivo*, establishing that translation from the IAV genomic strand can occur *in vivo*.

### Immunosurveillance Extends to Peptides Inserted Elsewhere in Genomic RNA

To extend these findings, we inserted SIINFEKL-encoding nucleotides elsewhere in genomic RNA. To minimize the perturbation to coding sequences, we BLASTED reverse strand ORF amino acid sequences against SIINFEKL to find the most homologous naturally encoded sequence and then changed the negative strand sequence to encode SIINFEKL. Unfortunately, inserting SIINFEKL at a position equivalent to residue 154 in HA (H3 numbering) or residue 809 of PA, which in each case changed only 4 residues, was incompatible with recovering infectious viruses.

We were, however, able to generate a virus (NArev_SIIN_) encoding negative strand SIINFEKL inserted in a position corresponding with residue 42 in the NA stalk (**Supplemental Fig. 4**). NA stalk length is naturally variable in nature, and NA is able to accept SIINFEKL in the positive strand with little consequence (31, 32). Indeed, NArev_SIIN_ grew to *wt* titers. SIINFEKL synthesis could potentially initiate at an AUG three codons upstream and would be terminated by a stop codon after translating 5 additional residues. The next potential start site is located dozens of codons upstream, and is unlikely to be relevant, since there are 6 intervening stop codons.

Although we failed to detect OT-I T cell activation with NArev_SIIN_ infected DC2.4 cells *in vitro* (**Fig. 2**), we observed weak but consistent OT-I T cell activation *in vivo* following i.p. infection, as assessed either by splenic and PEC OT-I T cell proliferation or IFN-γ production *ex vivo* (**Fig. 4**).

Thus, even in the absence of any selective pressure for gene expression as might occur with NEG8 SIINFEKL, there is sufficient translation from essentially a random sequence in genomic IAV RNA to enable CD8+ T cell immunosurveillance *in vivo.*

## Discussion

Despite lurking for decades in the published IAV segment 8 sequence (35, 36) in full view of clever students of IAV, evidence supporting the translation of the NEG8 ORF in the context of a *bona fide* influenza infection is lacking. Using multiple antibody-based approaches to detect NEG8 (raising rabbit antibodies to predicted NH_2_- and COOH-terminal peptides, generating NEG8 fused genetically to antibody epitope tags (Myc, HA, SIINFEKL itself using a SIINFEKL specific mAb (37)), we failed to detect NEG8 in PR8-infected HeLa or MDCK cells by immunofluorescence or immunoblotting using extracts prepared by immersing cells in boiling SDS extraction buffer to maximize protein solubilization. It is likely that if the protein is synthesized in these cells, it is only in minute quantities.

Given the unusual nature of its “mRNA” it would not be surprising if NEG8 is only expressed physiologically under special circumstances *in vivo*. Perhaps expression is limited to a subset of the wide variety of cells types that can be infected by IAV, which include epithelial, endothelial, and hematopoietic cells. It is clearly of interest in future experiments to examine possible NEG8 translation in whole organ extracts and frozen sections of lungs and immune tissues of infected animals.

Our findings show that ribosomes in cultured cells and *in vivo* can initiate downstream of the first AUG on the NEG8 ORF and continue translating to the predicted COOH-terminus and the appended model CD8+ T cell peptide SIINFEKL. Such translation, may however, be exclusively related to immunosurveillance, in which case the translation products may be truncated and degraded rapidly, with encoded peptides being rescued from typically rapid destruction by binding to MHC class I, and potentially, class II molecules as well (38).

How might negative strand IAV RNA be translated? vRNA is synthesized from an intermediate form of RNA (complementary RNA) dedicated to vRNA synthesis (39). In theory, negative sense RNA should not be capped, polyadenylated, or exported from the nucleus, except for incorporation into budding virions as NP-coated genomic segments. These events may however occur at a level with functional consequences, but still below the radar of current methodologies. Further, none of these events are absolutely required for translation, particularly for viruses. There are many examples of cap- and polyA-independent mRNA translation, including the translation of peptides embedded in introns, which appears to occur in the nucleus (19). Viruses are virtuosos at manipulating the rules of translation, typically exploiting non-canonical translation while shutting down canonical translation to monopolize ribosomes and also manipulate innate host anti-viral responses (40).

The nuclear localization of IAV transcription may be key to negative strand translation. We previously reported that upon drug blockade of NA mRNA export from the nucleus of cells infected with NA-SIINFEKL PR8, NA protein expression is reduced to a far greater extent than K^b^-SIINFEKL generation (18), consistent with peptide generation from nuclear translation of the NA positive strand. It would be interesting to determine if NEG8_SIIN_ is translated/immunogenic in the context of a typical cytoplasmic negative stranded virus such as VSV. If so, this would undermine the potential contribution of nuclear translation or aberrant splicing of genomic and mRNA to the SIINFEKL synthesis in NEG8_SIIN_ infected cells.

In summary, we show that, through an unknown mechanism, immunosurveillance extends to the negative strand of at least a subset of RNA viruses. If this is a general phenomenon, it implies negative strand information can potentially double the peptides presented for T immunosurveillance of viruses and cancer cells. If such T cells exert sufficient anti-viral activity to limit transmission, they could select for escape mutants that will contribute to viral evolution, and whose significance will not be apparent from standard analysis of viral ORFs.

## Material and Methods

### Confocal Microscopy

Hela cells were cultured on Nunc Lab-Tek chambered coverglass (Thermo Fisher Scientific, Waltham MA) for 24 h in normal growth media. Cells were infected with rVV-NEG8-GFP for 12 hours, fixed with cold acetone and incubated with rabbit polyclonal Abs specific for TGN46 (NB110-40769, Novus Biologicals, Littleton CO) and mouse PDI (ab2792, Abcam, Cambridge MA) following by Alexa 594 conjugated anti-rabbit and Alexa 647 conjugated anti-mouse IgG antibodies (Jackson ImmunoResearch, West Grove PA). Counterstaining was performed with Hoechst 33258 (Thermo Fisher Scientific, Waltham MA). Stained cells were visualized by SP8 confocal microscope system (Leica Microsystems, Mannheim Germany) using 405, 488, 561 and 633 nm excitation wavelength for blue, green, red and far red channels, respectively.

### *In vitro* OT-I activation

Animal work was performed with the approval of the NIAID animal care and use committee. Spleen and LNs were removed from OT-I TCR transgenic dsRed mice and homogenized to produce single-cell suspensions. Red blood cells were lysed, and samples were filtered through a 70-µm nylon filter, and cells purified using an AutoMacs and the CD8+ T cell Negative Selection Kit (Miltenyi Biotech) according to the manufacturer’s instruction. Purified cells were labeled in PBS with the Cell Trace Violet Cell Proliferation Kit (ThermoFisher) according to the manufacturer’s recommendations. Cells were plated at 5 × 10^4^ cells/well in 96-well U-bottom plate. DC2.4 cells were infected with at an MOI ~ 10 for approximately 1 hour, then added at 1 × 10^5^ /well. Some wells received uninfected DC2.4 cells pulsed with 0.1 µm SIINFEKL peptide as a positive control. Percent IAV infection was determined 8 h PI via flow cytometry for cell surface HA expression. Twenty-four and forty-eight h post-infection, cells were stained for CD8 (53–6.7), CD69 (H1.2F3), and CD25 (PC61) (all from eBioscience) and analyzed for Cell Trace Violet fluorescence to determine cell division on a BD LSRII flow cytometer (BD). Results were analyzed using FlowJo (Tree Star).

### *In vivo* activation of OT-I cells

Twelve to 24 h prior to infection, we transferred 2 × 10^5^ CD45.1^+^ OT-I cells (purified as above) i.v. into C57Bl/6 (CD45.2^+^) mice. For analysis of IFN-γ production, mice were infected i.p. with ~1 × 10^8^ TCID50 of each recombinant. Spleens or peritoneal exudate cells (PECs) were harvested at 7 days PI, homogenized, and cells resuspended in RPMI-10 + 10 mM Hepes buffer and plated at 2 × 10^6^ cells/well in U-bottom, 96-well plates along with SIINFEKL or an irrelevant control peptide (SSIEFARL) at a final concentration of 100 nM. Cells and peptide were incubated for 4 h. at 37°C in the presence of 10 µg/ml brefeldin A (Sigma-Aldrich) to allow IFN-γ accumulation. For dead cell exclusion, cells were treated with ethidium monoazide (Invitrogen) before washing, incubating with Fc block (antibody 2.4G2 produced in house), and staining with anti-CD8 (clone 53–6.7) and anti-CD45.1 (clone A20; eBioscience). After staining, cells were washed and fixed at room temperature for 15 min with 1% paraformaldehyde. Cells were incubated overnight at 4°C with Alexa Fluor 647 anti–IFN-γ (clone XMG1.2, eBioscience) diluted in PBS containing 0.5% saponin (EMD Biosciences). Cells were analyzed on a BD LSR II and results analyzed using FlowJo.

### Peptide Binding Assay

Peptide binding was determined as described (41). Briefly, highly purified synthetic peptides were dissolved at 1 mM in DMSO and diluted in FBS-free DMEM to limit proteolysis. RMA/S cells cultured overnight at 27°C, washed, were then incubated with indicated concentrations of peptides for 2 h at room temperature, incubated in the presence of 5 μg/ ml brefeldin A at 37°C for 2 h, and then washed and stained with anti-Kb antibody AF6-88.5. Secondary staining was conducted with Alexa Fluor 647-coupled goat anti-mouse IgG(H+L) (Life Technologies).

### Plasmid Mutagenesis

Mutant Influenza A viruses created were all based on the Mt. Sinai PR8 strain molecular clone. Plasmid pDZ-PR8-NS was mutagenized by oligonucleotide-based site-directed mutagenesis using the QuikChange system (Agilent Technologies, Waldbronn, Germany) according to the manufacturer’s instructions. Oligonucleotide sequences used for appending SIINFEKL to NEG8 were TTTCGAAGTTGATGATCGACCTAACTGACATGACTCTTGAG and CGATCATCAACTTCGAAAAGCTATAACGCGACGCAGGTACAGAGG; for appending a stop codon and SIINFEKL to NEG8 were TTTCGAAGTTGATGATCGATTACCTAACTGACATGACTCTTG and CGATCATCAACTTCGAAAAGCTATAACGCGACGCAGGTAC; for appending a stop codon, ACG codon (Thr) and SIINFEKL to NEG8 were TTTCGAAGTTGATGATCGACGTTTACCTAACTGACATGACTCTTG and CGATCATCAACTTCGAAAAGCTATAACGCGACGCAGGTACAGAG; for mutagenizing the predicted AUG start codon of NEG8 to ACG were GAATAGTTTTGAGCAAATAACgTTTATGCAAGCCTTACATC and CTTATCAAAACTCGTTTATTGcAAATACGTTCGGAATGTAG; for appending the myc tag to the predicted C-terminus of NEG8 were ACAGGTCCTCCTCGGAGATGAGCTTCTGCTCCCTAACTGACATGACTCTTG and CATCTCCGAGGAGGACCTGTAACGCGACGCAGGTACAGAGG; and for appending three tandem myc tags to the predicted C-terminus of NEG8 were CTTCTTCTGAAATCAACTTTTGTTCCAGATCTTCTTCAGAGATGAGTTTCTGCTCtCCtCCCCTAACTGACATGACTCTTG and TGATTTCAGAAGAAGATCTGGAACAGAAGCTCATCTCTGAGGAAGATCTGTTAATTAATTGACGCGACGCAGGTACAGAG.

### Virus Rescue and Propagation

Recombinant viruses were rescued using an <u>eight plasmid system.</u> One microgram of each of the eight wild-type and mutant plasmids and two micrograms of pDZ-PR8-NP were mixed in 186 µl Optimem medium (Life Technologies). 16 µl of Trans-It LT1 (Mirus Bio, Madison WI) was added and mixed, followed by incubation at room temperature for one hour. 800ul Optimem was added and the mixture pipetted onto an aspirated 6-well well of cells plated the previous day at 0.6 × 106 293-T cells plus 0.2 × 10^6^ MDCK cells in DMEM with 7.5% fetal calf serum. Transfected cells were incubated overnight at 37°C.

The following morning, medium was replaced with 3 ml DMEM without serum containing 50 µg/ml gentamycin (Quality Biological, Gaithersburg MD), 3 µg/ml L-(tosylamido-2-phenyl) ethyl chloromethyl ketone (TPCK)-treated trypsin (Worthington Biochemical, Lakewood NJ) and 25 mM HEPES (Quality Biological). Over the following days, supernatant media was removed and assayed for hemagglutinin activity. To grow virus further, 50 µl rescued virus supernatant was injected into incubated 10-d embryonated pathogen-free hen eggs (Charles River, Norwich CT). 48 h post-infection, allantoic fluid containing the virus was removed and stored at −80°C.

## Acknowledgements

This work was supported by the Division of Intramural Research, NIAID.

---

**Supplemental Figure 1.**
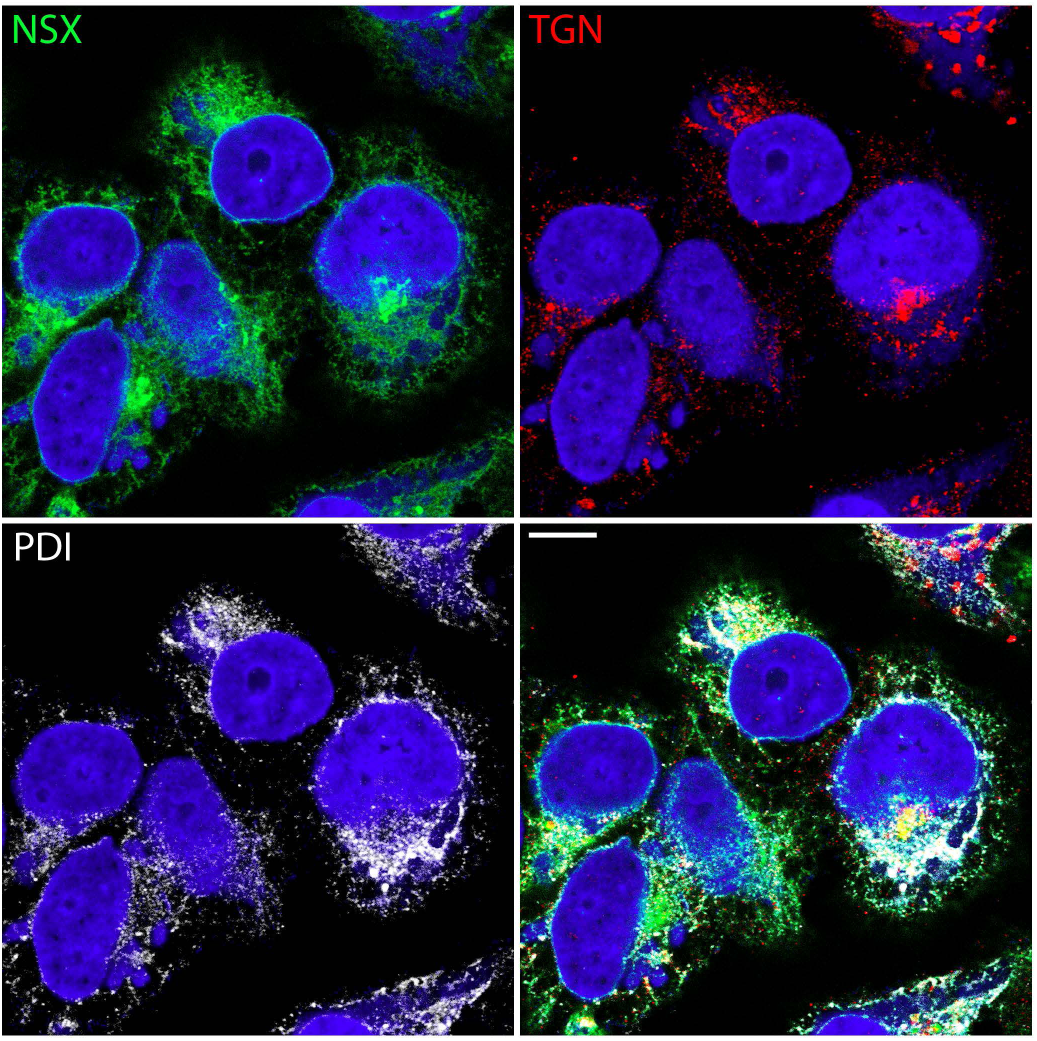
rVACV-expressed Neg8GFP localizes to the ER and GC. HeLa cells infected with a rVACV expression Neg8GFP for 12 hours were stained for ER (white) and GC (red) markers and nuclei and viral factories (blue) and imaged by confocal microscopy. Lower right panel merges the individual color panels. Scale bar is 10 microns. The experiment was repeated 3 times with similar results.

**Supplemental Figure 2.**
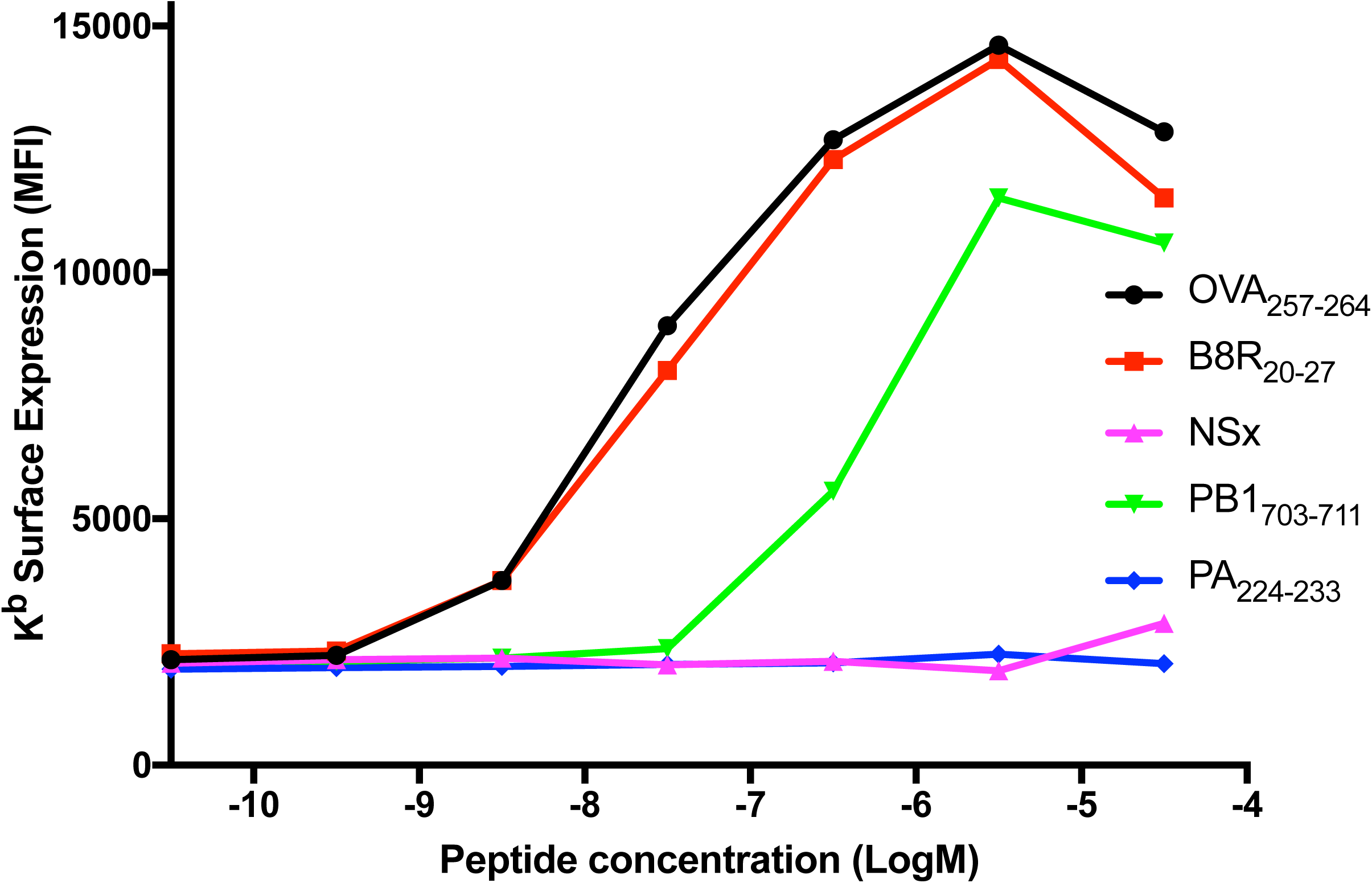
Previously Identified Neg8 Peptide Does Not Bind K^b^. RMA/S cells (TAP-deficient (42)) were cultured overnight at 27 °C to accumulate peptide receptive K^b^ molecules. Cells were then incubated with peptides at the indicated concentration at room temperature and then shifted to 37 °C for 2 h to denature peptide free K^b^ molecules. Cells were then stained on ice with directly conjugated anti-K^b^ mAb to determine peptide bindings. B8R and OVA (SIINFEKL) peptides bind K^b^ with high affinity, while PB1 binds with intermediate affinity. PA is a D^b^ binding negative control peptide. The NSX peptide tested exhibits no significant binding to K^b^. The experiment was repeated 3 times with highly similar results.

**Supplemental Figure 3.**
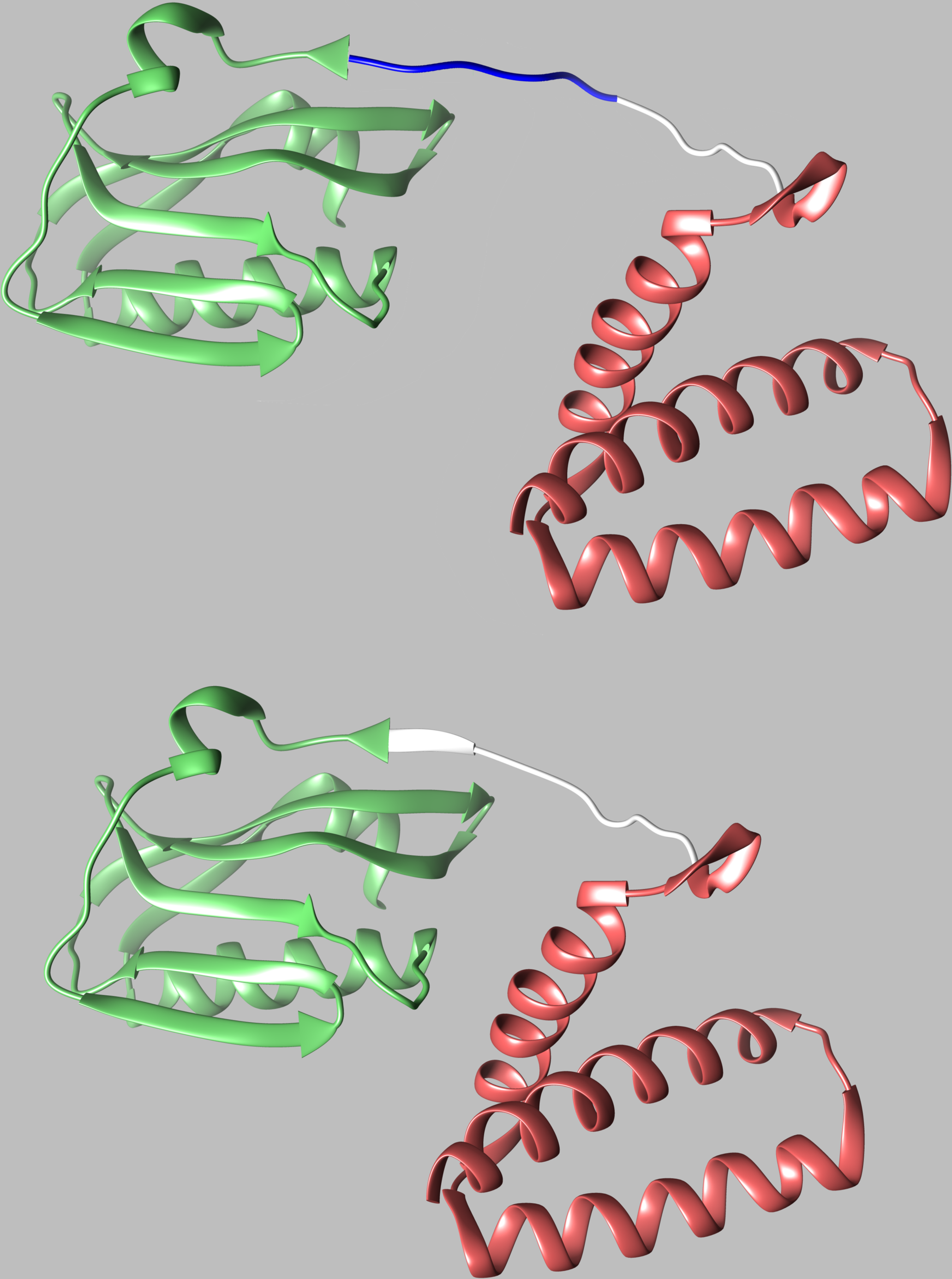
Insertion of eight amino acid peptide into linker sequence of NS1. Appending SIINFEKL to the carboxy-terminus of NEG8 results in the insertion of the sequence SFSKLMID (shown in blue) after tyrosine 89 of the linker (amino acids 79–89, shown in white) between the RNA-binding (red) and effector (light green) domains of NS1. Molecular graphics and analyses were performed with the UCSF <u>Chimera</u> package (43) using PDB file 4OPH, “X-ray structure of full-length H6N6 NS1” retrieved from rcsb.org. Chimera is developed by the Resource for Biocomputing, Visualization, and Informatics at the University of California, San Francisco (supported by NIGMS P41-GM103311).

**Supplemental Figure 4.**
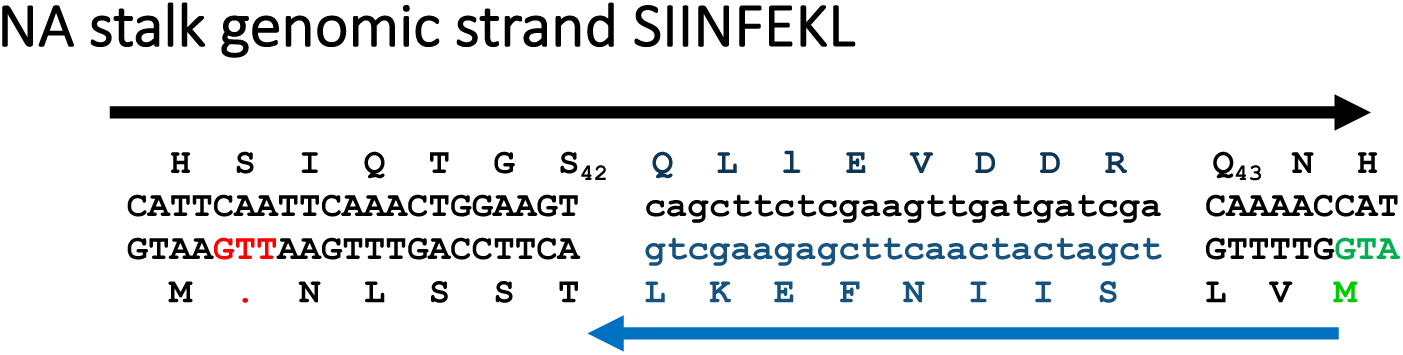
Construction of NArev_SIIN_ Virus. We inserted the sequence indicated (nucleotides in small letters) encoding SIINFEKL in the negative sense of segment 6 RNA resulting in the complementary peptide (QLLEVDDR) inserted into the NA stalk between residues 42 and 43. SIINFEKEL (blue) can be initiated by the Met indicated in green, and a translating ribosome would stop (red dot) after translating 5 additional residues (TSSLN). Upstream of the indicated AUG are 6 stop codons prior to the next AUG, making it a very unlikely initiation codon for SIINFEKL translation.

